# Systemic and vascular inflammation in an in-vitro model of central obesity

**DOI:** 10.1101/188391

**Authors:** A. Ahluwalia, A. Misto, F. Vozzi, C. Magliaro, G. Mattei, MC. Marescotti, A. Avogaro, E. Iori

## Abstract

Metabolic disorders due to over-nutrition are a major global health problem, often associated with obesity and related morbidities. Obesity is peculiar to humans, as it is associated with lifestyle and diet, and so difficult to reproduce in animal models.

Here we describe a model of human central adiposity based on a 3-tissue system consisting of a series of interconnected fluidic modules. Given the causal link between obesity and systemic inflammation, we focused primarily on pro-inflammatory markers, examining the similarities and differences between the 3-tissue model and evidence from human studies in the literature. When challenged with high levels of adiposity, the in-vitro system manifests cardiovascular stress through expression of E-selectin and von Willebrand factor as well as systemic inflammation (expressing IL-6 and MCP-1) as observed in humans. Interestingly, most of the responses are dependent on the synergic interaction between adiposity and the presence of multiple tissue types. The set-up has the potential to reduce animal experiments in obesity research and may help unravel specific cellular mechanisms which underlie tissue response to nutritional overload.

## 1. Introduction

Overweight and obesity are major risk factors for a number of chronic diseases, including diabetes [1], cardiovascular diseases and cancer [2]. Since the turn of the century, the number of obese adults has increased to over 300 million. Obese individuals often have excess central visceral adiposity, a condition that contributes to a chronic increase in circulating free fatty acids and metabolites, such as glycerol and triglycerides. These metabolites in turn activate various signaling cascades that interfere with insulin signaling and β-cell function, further contributing to gluco/lipotoxicity [3].

A great deal of research has been dedicated to delineate the etiopathogenic mechanisms of obesity and diabetes using animal models. The most widely employed models of obesity are rodents, either mutant or genetically engineered mice or rats in which adiposity is induced by prolonged feeding on high fat diets [4,5]. As succinctly put by Wang et al, “*despite the extensive use of these rodent models, many mechanistic details of human metabolism remain poorly understood and treatment options for humans are limited and largely unsatisfactory*” [6]. Indeed, we now know much about the details of rodent metabolism, but still lack a detailed understanding of the mechanisms underlying human glucose homeostasis and response to chronic over-nutrition as well as human obesity related comorbidities and responses to interventions [7].

Besides the evident differences between human and rodent lifespan, diet, and basic biology, animal models are not amenable to dissociation of metabolite dynamics in different tissues and organs. As a consequence, identifying the contribution of individual tissues or organs to nutrient balance or destabilization is a formidable task. Some attempts have been made to model human obesity using in-vitro techniques, most of these deal with adipose cell or tissue cultures, derived from cell lines or isolated from donors [8,9]. In-vitro research has contributed significantly to our understanding of alterations in signaling at the membrane or cytoplasmic level in single cells [10,11]. Thus, although there is clearly a network of signaling between different tissues which contributes to maintain energy homeostasis in the human body, much of our understanding of signal transduction is limited to a very small space and time window. The question of how metabolic signals are propagated to and translated by remote tissues and organs, and how the internal milieu is modulated by them has not yet been addressed, with very little research carried out on higher level models of endogenous metabolism containing different cell or tissue types.

Several microfluidic systems involving multiple tissues are now emerging, while most are focused on systemic toxicity [12,13], Xiao et al recently demonstrated that reproductive tract tissues can be cultured together under flow to produce a semblance of the human menstrual cycle hormonal profile [14]. However, as far as nutrient metabolism is concerned, very little work has been done on models involving the interactions between different cells or tissues linked in the body through the vascular network. In fact, the metabolic circuitry involves all organs and tissues, most of them as end receivers of energetic substrates, which in normal resting conditions, are generally glucose and some free fatty acids (FFAs) [15]. Some organs and tissues play more important roles such as the engineering equivalents of processing, sorting, storage and control [16]. Recapitulating systemic metabolism is a challenging task and requires a methodical reverse engineering approach, breaking down the circuitry to its most basic elements and reconstructing the metabolic network in-vitro. By increasing the number of interactions and variables step-wise in a properly scaled model, nutrient dynamics between organs and their contribution to the whole body metabolic profile can be systematically investigated [17].

Using this approach we previously reported a modular in-vitro system for the study of glucose and lipid dynamics in a model of central metabolism [18,19]. The system consists of interconnected bioreactor chambers linked together by the flow of a common medium, containing respectively human adipose tissue, human endothelial cells and a human hepatocyte cell line. Cell ratios were scaled proportionally to represent adipose tissue, vascular tissue and liver in the abdomen [20,21]. This paper builds on a series of studies in which we demonstrated that fluid flow reduces glucose uptake and increases lactate availability in all cells, particularly hepatocytes and endothelial cells [22]. Hepatocytes are the master regulators in the 3 tissue fluidic connected culture (herein referred to as 3-way), as their introduction to the 2 tissue (2-way) endothelial cell-adipose tissue circuit markedly reduced changes in metabolite concentrations with respect to fresh media levels [18]. Following these initial investigations, the 3-way system was challenged with high glucose concentrations with and without insulin to simulate type I and type 2 diabetes respectively. The results demonstrated that the system’s metabolic and inflammatory profile changes dramatically in the presence of high concentrations of glucose, and that these changes are modulated by the presence of insulin, highlighting its potential to recapitulate some of the characteristics of human metabolism in the presence of nutritional overload [23]. Here, the 3-way model was refined to include 3D porous collagen scaffolds, a high shear vascular compartment and different levels of human adiposity, representing normo-weight with 12% adipose tissue (12%AT), overweight (25%AT) and obese (35%AT) conditions respectively. The aim of the study was to challenge the in-vitro system with increasing levels of adiposity to determine the extent to which the 3-tissue fluidic system reflects salient features of obesity-related vascular and systemic stress observed in humans.

## 2. Material and methods

### 2.1. Cells

As in our previous studies, cell proportions in the 3-way set up represented the hepatocyte:adipocyte:endothelial ratio in the visceral region, i.e. 12%AT 10:2:1 for normoweight; 25%AT 10:4:1 for overweight; 35%AT 10:6:1 for obese [18].

Human Umbilical Vein Endothelial Cells (HUVEC, passage 4 to 6) were from Promocell. They were cultured in ECGM (Endothelial Cell Growth Medium, Promocell) with 10% di FBS (Fetal Bovine Serum), 0.1 ng/mL EGF (Epidermal Growth Factor), 1.0 ng/mL BFGF (Basic Fibroblast Growth Factor), 90 μg/mL heparin, 1.0 μg/mL hydrocortisone, and pen-strep. This cocktail is henceforth referred to as the common media. The cells (38 000) were pipetted into the μ-Slides which had been prepared by coating with 200 μL of 1% gelatin followed by 24 h pre-incubation in a 37°C oven.

The hepatocyte cell line HepG2 was from ATCC (American Type Culture Collection). These cells maintain the principal endogenous functions of human hepatocytes and respond to glucose-6-phosphate, conserving their capacity to synthesize glycogen [24]. Hepatocytes were passaged in EMEM with 1g/L glucose, 5% FBS and pen-strep. 150,000 cells were carefully pipetted into the center of porous collagen scaffolds (porosity 98%, pore size 200 μm, elastic modulus 1.2 kPa) placed in 48-well plates. The cells were incubated in the common media 48 hours before starting the connected culture experiments. After this time the cells proliferate to about 400,000-450,000, as ascertained by cytometry, and are ready for the 3-way experiments.

Visceral adipose tissue was obtained from n=9 donors undergoing liver resection for metastatic/benign liver lesions with no underlying chronic liver disease or diabetic complications. All patients provided their informed consent to participate in the study, which was approved by the Local Ethical Committee (Study n. 3059 approved on 21/07/2011 by the Azienda Ospedaliera Università Pisana). The study was carried out in accordance with guidelines established by the Local Ethical Committee. The tissue was cleaned of visible blood vessels and divided into weighed portions representing i) normoweight, 56 mg, ii) overweight, 112 mg, and iii) obese, 168 mg. The weighed samples were placed in wells and partially digested in collagenase (Sigma) for 10 min. They were then rinsed in EMEM with 20% FBS and pen-strep before inserting in the bioreactors. Using this procedure, we obtain a yield of about 1500 adipocytes/mg [22].

### 2.2. Bioreactors

The bioreactors were LB1 and LB2 (IVTech srl IT) and μ-Slide (Ibidi DE). LB 1 is a 24-well sized transparent mill-scaled chamber for fluidic culture of scaffolds and membranes under low shear stress, while LB2 has 2 flow inputs and outputs (Figure 1). The 400 μm channel height μ-Slide is designed for simulating the high shear vascular environment.

**Figure 1:**
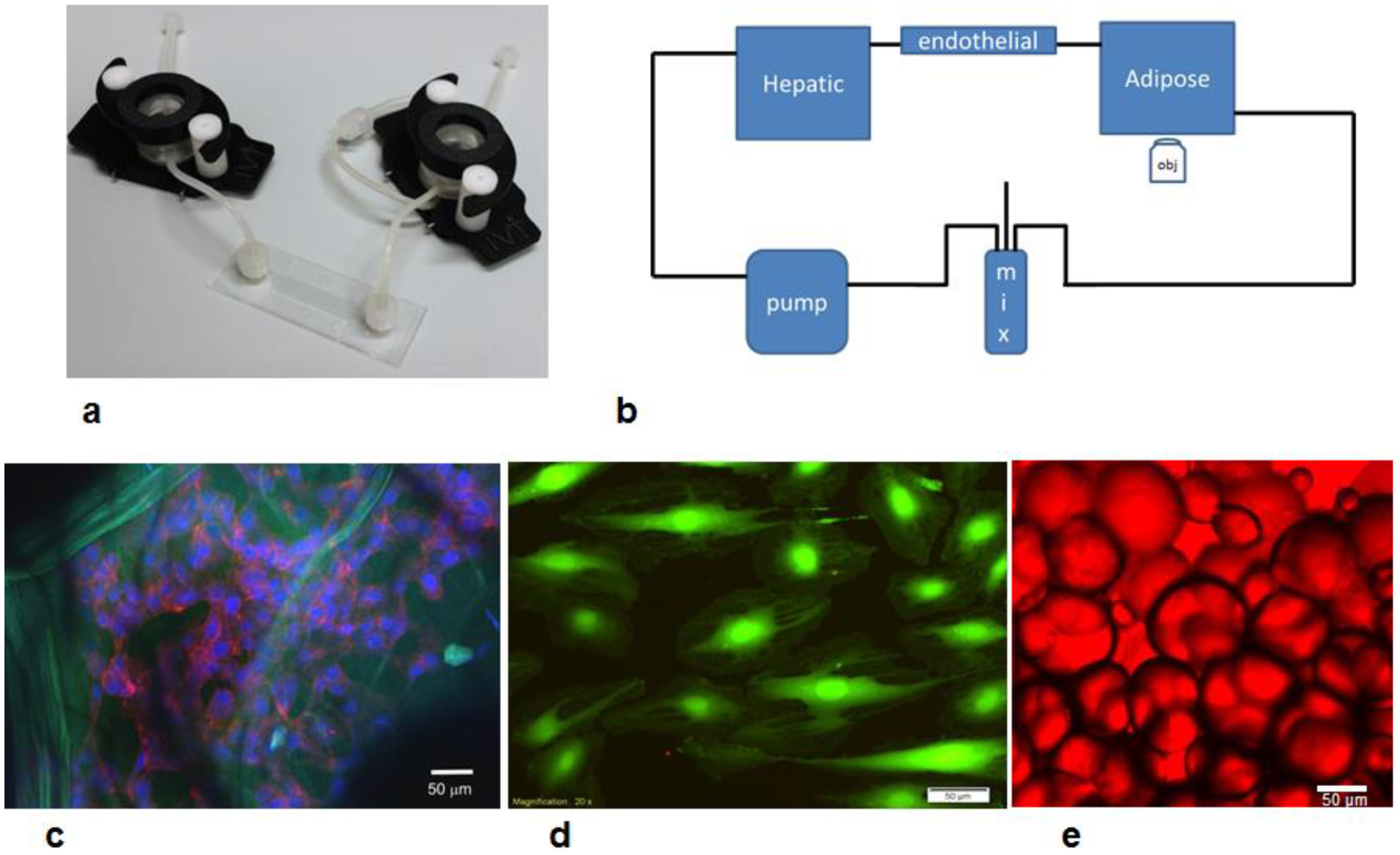
a) Basic components of the 3-way fluidic system: the LB1 chamber, the Ibidi laminar flow chamber, and the LB2 chamber; b) the fluidic set up with a peristaltic pump, mixing chamber and hepatic (LB1), endothelial (Ibidi) and adipose tissue (LB2) chambers; c) HepG2 cells stained with DAPI (blue) and actin-phalloidin (red) seeded on porous collagen scaffolds (green autofluoresence). Scale bar 50 μm; d) endothelial cells stained with calcein. Scale bar 50 μm; e) adipocytes stained with oil-red. Scale bar 50 μm.

An LB2 chamber was used for adipose tissue employing a cross-flow configuration for non-adherent cultures as recommended by the manufacturer. The tissue was contained in the chamber by piece of sterile nylon mesh. Hepatocytes seeded on collagen sponges were placed in an LB1 chamber while the μ-Slide was used for endothelial cells. In the 1-way experiments, each cell type was placed in the flow circuit singly. The 2-way experiments were conducted by connecting an LB2 chamber with adipose tissue to a μ-Slide with HUVEC. Finally, in the 3-way tests, one LB1 chamber with hepatocytes was added to the 2-way circuit. In all experiments (i.e. 1-way, 2-way and 3-way, see below), the total circuit volume was 14 mL, and the common medium was complete ECGM, which conserves the vitality of HepG2 and adipose tissue with respect to their standard media [22].

After seeding, the chambers were assembled using Luer connectors to form a closed loop circuit containing a peristaltic pump (Ismatech, CH) and a mixing chamber (IVTech srl, IT) as shown in Figure 1. The flow rate was 250 μL/min which gives a wall shear stress of 5 μPa in LB1 [25], and 0.35 Pa in the μ-Slide.

### 2.3. 1, 2 and 3-way cultures

The metabolic and inflammatory response of normo-weight, overweight, and obese (respectively 12%AT, 25%AT and 35%AT) amounts of adipose tissue was investigated in 1-, 2- and 3-way cultures using transparent bioreactors linked in series (Figure 1a, b). Firstly, using the same volume of ECGM medium in all tests, the fat compartment was characterised to obtain internal controls to account for variations due only to the mass of adipose tissue in a 1-way AT system. One-way responses of hepatic and endothelial cultures in the fluidic system were also assessed. Then an intermediate 2-way circuit with endothelial cells and adipose tissue was established, followed by a 3-way circuit with hepatocytes seeded on porous 3-dimensional collagen scaffolds, endothelial cells and adipose tissue. Different percentages of adipose tissue were added to the 2- and 3-way system in order to mimic different degrees of adiposity, keeping the hepatic and endothelial compartments constant. The metabolic profile and pro-inflammatory responses were evaluated to clarify the role of adiposity and the contribution of the different cell types in modulating media metabolite and stress-marker levels. Glucose, human albumin, urea and lactate levels were also monitored over time. As illustrated in Fig 1c-e, the cells were vital and adipocyte dimensions were conserved over the 24 hours of the experiments.

### 2.4. Cell viability

The viability of all three cell types in the connected cultures was measured and evaluated through the lactate dehydrogenase (LDH) assay using the method described by Decker and Lohmann-Matthes [26].

### 2.5. Biomolecule analyses

In order to assess the cell functions and viability, 200 μL of media were withdrawn from the mixing chamber at fixed intervals (0, 4, and 24 h). Biochemical assays were performed to measure the change over time in the metabolite and pro-inflammatory marker levels in the media.

Free Fatty Acids (FFAs) and triglycerides were measured by a colorimetric enzyme assays (respectively NEFA C test-Wako Chemicals GmbH, Germany and Hagen Diagnostica SRL, S. Giovanni V.no (AR), ITALY). Glycerol was determined by a modified Lloyd assay using an automated spectrophotometer Cobas Fara II (Roche) [27]. IL-6, (eBioscience Dx Diagnostic, Vienna, Austria), E-Selectin, (Boster Biological Technology, LDT, Tema Ricerca, Bologna, Italy), MCP-1 (Life Technologies Italia, Monza-MI, Italy, albumin (Bethyl Laboratories, Montgomery, TX, USA), were determined using ELISA. Other assays and their results are reported in the SI.

At the end of the experiments the cells were fixed and stained with DAPI and phallodin or anti-vWF (all from Thermo Fischer, Italy) and observed with a confocal (Nikon A1, IT) or fluorescent microscope (Olympus X81, IT). Adipose tissue was stained with oil-red stain. The intensity of vWF staining was quantified by image processing as described in the SI.

### 2.6. Data Analysis

Results are reported as the mean ± standard deviation, unless otherwise noted. The statistical differences in metabolite and cytokine concentrations (with respect to fresh media) between the three levels of adiposity in 1-way, 2- and 3-way connected cultures was analysed using two-way ANOVA followed by Tukey’s Multiple Comparison Test in order to evaluate both the effect of varying the adiposity for a given model complexity (i.e. same way) and that of changing model complexity for a given level of adiposity. Since human albumin is only secreted by the hepatocytes (as the medium contains bovine albumin), its production was evaluated only for the most complex 3-way hepatocyte-containing circuit: the effect of adiposity in the 3-way system was tested with 1-way ANOVA followed by Tukey’s Multiple Comparison Test, and results compared to the basal albumin production of the hepatocyte monoculture. Similarly, statistical significance between selectin and vWF levels for the different 3-way conditions and the single 1-way controls was determined with one-way ANOVA. Post-hoc multiple comparisons between different groups of data were carried out using the Tukey’s test. Statistical analysis was implemented in Graphpad Prism 6.0. Differences were considered significant at p < 0.05. At least 3 different experiments were conducted for each condition, using adipose tissue from a total of 9 different donors. Further details on the image processing methods are reported in the Supplementary Information.

## 3. Results

### 3.1. Metabolic profiles

Figure 2 shows the average fractional change in key metabolites at 24h for different dynamic conditions. The star charts are useful for rapidly assessing how the addition of different tissues to the fluidic circuit changes the overall metabolic fingerprint of the system. Metabolite levels are expressed as the ratio of the change in concentration with respect to fresh medium. The metabolic profiles of 1-way HUVEC and hepatocytes in Figure 2a are similar to our previously published data [22] and indicate that HUVEC release FFA into the medium while hepatocytes are a source of human albumin. Figure 2b and c show peaks in lactate, glycerol and FFA levels which are not correlated with the amount of adipose tissues and are probably due to donoro-to-donor variations in human adipose samples. The star plot in Figure 2d shows that media concentrations are similar for all metabolites in the 3-way circuit except for urea and lactate levels in the presence of the highest percentage of AT. Thus, the addition of hepatocytes to the 2-way circuit reduces the differences in metabolite levels between the different adipose tissue conditions. This ‘regulatory’ effect of hepatocytes was also observed in in our previous studies [28].

**Figure 2:**
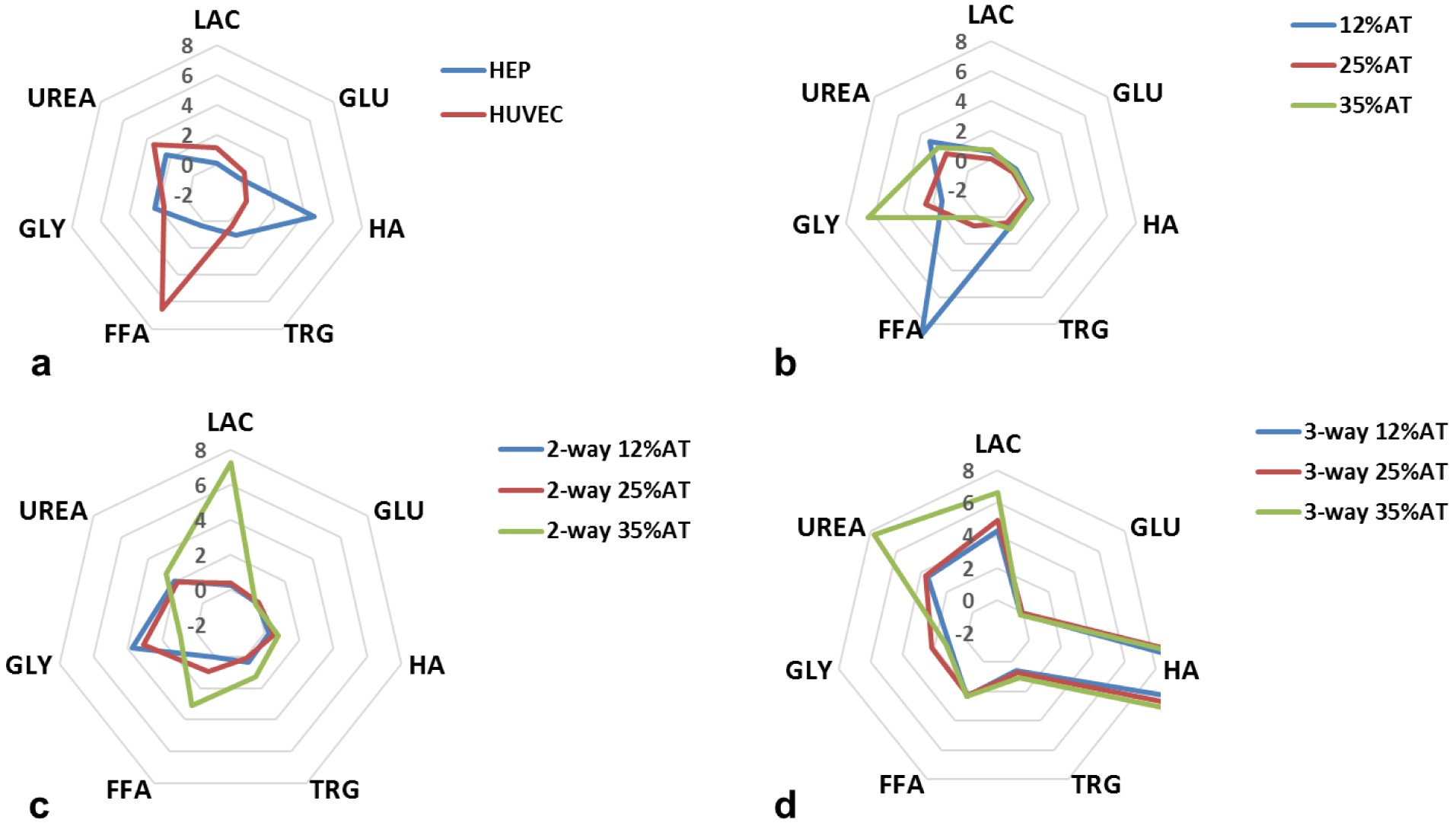
Metabolic profiles: a) 1-way hepatocyte and HUVEC; b) different quantities of 1-way adipose tissue (AT); c) 2-way HUVEC with increasing quantities of AT; d) 3-way connection with increasing quantities of AT (the data point for HA is not on the chart as it increases over 20-fold in the 3-way circuit). The data refer to average fractional changes in metabolite concentration after 24h of culture in the bioreactor system with respect to fresh media levels. LAC: lactate, GLU: glucose, HA: human albumin, TRG: triglycerides, FFA: free fatty acids, GLY: glycerol (n=3 independent experiments for data each point).

### 3.2. Lipids and albumin

The effect of both adiposity (i.e. 12, 25 and 35 %AT) and model complexity (i.e. 1-, 2- and 3-way) on the production of triglycerides (TRG), glycerol (GLY), FFAs and albumin (HA) after 24 h of culture was investigated in order to determine whether and how lipids and lipid-related metabolite levels in the system are modulated by the amount of adipose tissue considered in the model and/or by cross-talk between multiple tissues. Significant interaction between adiposity and the level of cross-talk was found for TRG production (p = 0.0039), meaning that the effect of the first factor is dependent on the level of the second one, and vice-versa. Overall more triglycerides were released from cells in the culture medium over 24 h in the 3-way cultures (Fig. 3a). In particular, 3-way TRG production was higher than 2-way at 12% adiposity (p=0.0138), while it was higher than both 1- and 2- way at 25% adiposity (p<0.0001). However, in neither case did the concentrations correlate with the percentage of adipose tissue in the system. Similarly, the two variability factors investigated (i.e. adiposity and system complexity) also interact significantly for glycerol production (p<0.0001). Glycerol concentrations were increased over 24 hours in all the 3-way circuits with respect to the 1-way adiposity controls, particularly in the 25%AT system where the levels were more than doubled with respect to all the other conditions (2-way ANOVA multiple comparison analysis p<0.037, Fig 3b). Significant variable interaction was also found for FFAs (p<0.0001). In particular, their production significantly decreases with increasing adiposity in the 1-way AT cultures (p<0.0003). On the other hand, although we observed a net release of FFAs in the 3-way media over 24 hours (Fig. 3c), the levels were very similar independent of the percentage of adipose tissue added to the system. In fact, the 2-way ANOVA interaction analysis shows that FFA levels depend on adiposity (p<0.0001), but only in the absence of other cell types.

**Figure 3:**
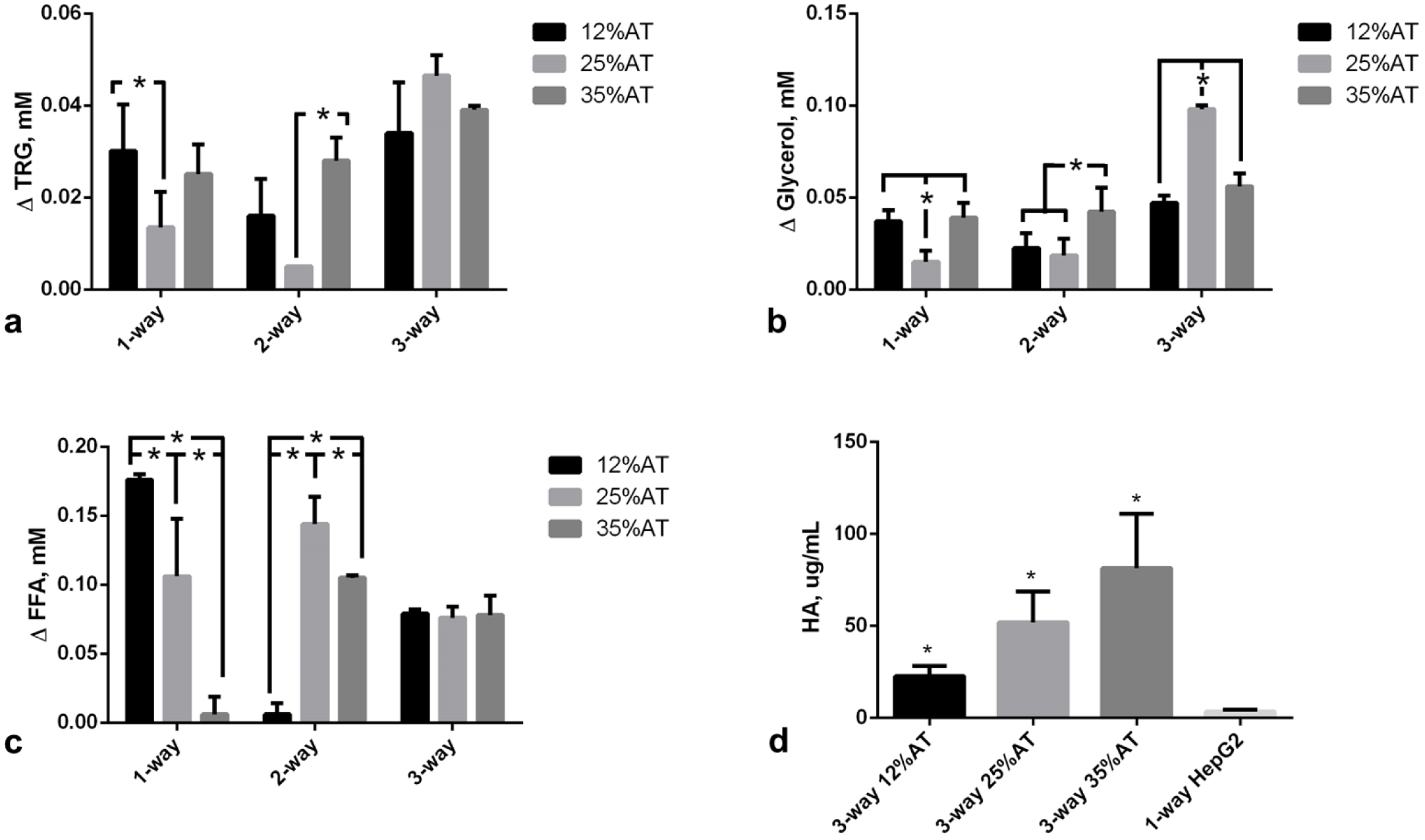
Change in metabolite concentrations with respect to fresh media after 24h. a) Triglycerides (TRG); b) Glycerol; c) FFAs; d) Human albumin (HA). For a), b), and c) the change AT content in 1-, 2- and 3-way cultures was compared within the same group, *= p<0.05. For d) the control is a 1-way hepatocyte culture. *= p<0.05 with respect to the corresponding 1-way control (n=3 independent experiments for data each point).

Albumin secretion increased with the amount of adipose tissue in the 3-way circuit (Figure 3d), being significantly higher at 35% adiposity with respect to 12% AT (p=0.0264). Hepatocytes seeded on collagen scaffolds in the absence of other cells in connection secreted significantly less albumin (3.38±1.28 μg/mL, p=0.004 with respect to 3-way 12%AT, 1-ANOVA) over 24 h. As albumin is a carrier of fatty acids, its increased secretion by hepatocytes may be directly associated with the increase in adiposity and correlated with the levelling of FFA concentrations observed in Figure 3c. These data suggest that the 3-tissue cross-talk stabilises media FFA concentrations, likely through increased triglyceride (Fig. 3a) and albumin (Fig. 3d) production by hepatocytes. Additional data on complete 2-way ANOVA analyses and intermediate time-points are reported in the Supplementary Information (SI).

### 3.3. Pro-inflammatory markers

Two systemic pro-inflammatory markers were investigated: Interleukin-6 (IL-6) and Monocyte chemoattractant protein-1 (MCP-1). Our data confirmed that there was a significant (p<0.001) increase in IL-6 when HUVECs and hepatocytes were connected to 35%AT or 25%AT, as shown in Fig. 4a. The synergic interaction between adiposity and connectivity giving rise to high levels of this cytokine in the presence of 25 and 35% adipocytes in the 3-way group was confirmed by the interaction analysis (p<0.0001, 2-way ANOVA) (Figure 4b). MCP-1 levels were highly correlated with adiposity in both the controls and 3-way condition, an indication of insignificant variable interaction (p=0.1637, 2-way ANOVA). In the media of 3- way connections with 25%AT we found a net increase in MCP-1 medium concentrations with respect to those observed in presence of 12%AT (p=0.0022) or 35%AT (p=0.0183), Figure 4b. IL-6 and MCP-1 were close to the detection limit in the 1-way HUVEC and 1-way hepatocyte circuits.

**Figure 4:**
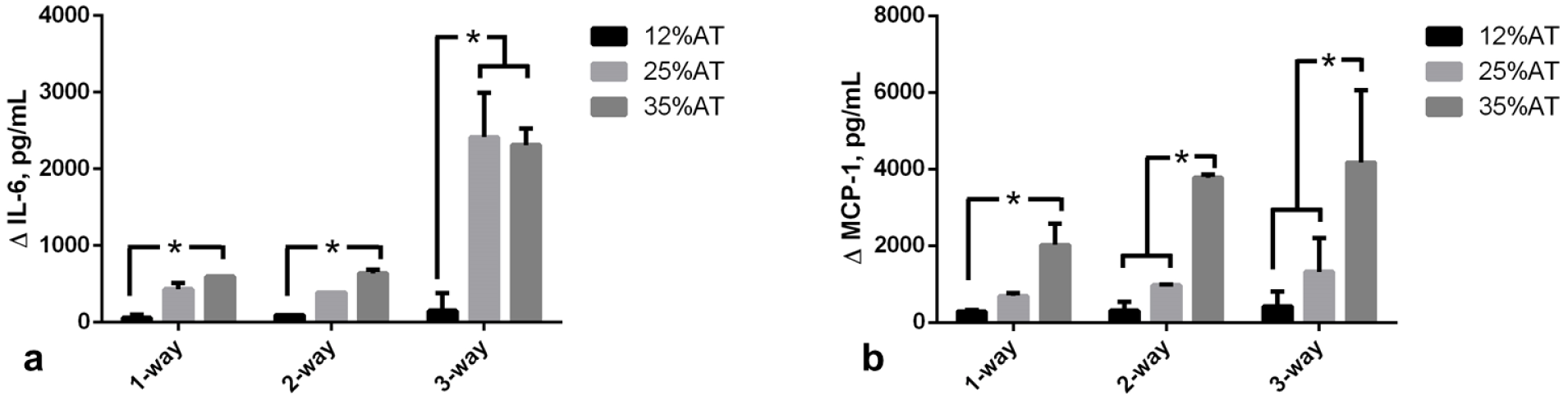
Pro-inflammatory markers at 24h. a) Changes in IL-6 concentration in 1-, 2- and and 3-way connected cultures as a function of adiposity; b) Changes in MCP-1 concentration in 1-, 2- and 3-way connected cultures as a function of adiposity. *= p<0.05 (n=3 independent experiments for data each point).

### 3.4. Endothelium specific markers

As obesity related cardiovascular complications are closely associated with endothelial dysfunction [29], we also focused on two markers of endothelial stress in the 3-way system, comparing responses with 1-way controls with only HUVEC. E-selectin, is a cell adhesion molecule expressed only on endothelial cells activated by cytokines and is an early indicator of endothelial damage. Von-Willebrand Factor (vWF) is produced constitutively in the Weibel-Palade bodies of the endothelium and is a more tardive marker of endothelial stress with respect to E-selectin [30,31].

When 12%AT was connected to HUVECs and HepG2 cells, medium E-selectin levels were stable over 24 hours, with levels similar to those of 1-way HUVEC (Fig. 5a). As adiposity in the system was increased we observed a significant release of E-selectin (p<0.0001 35%AT vs. 12%AT and the control; p<0.0001 25%AT vs. 12%AT and the control), showing that the presence of a higher amount of fat can induce endothelial stress (Fig. 5a). vWF expression in HUVEC was measured using immunostaining and image quantification as described in the SI. A significant (over 2-fold, p=0.0012) increase in vWF fluorescence per cell was observed in going from HUVEC in the 3-way connection with 12%AT to 35%AT and 25%AT. No difference was observed between negative controls (HUVEC only) and HUVEC in 3-way with 12%AT. LPS treated cells were used as positive controls (Figure 5b).

**Figure 5:**
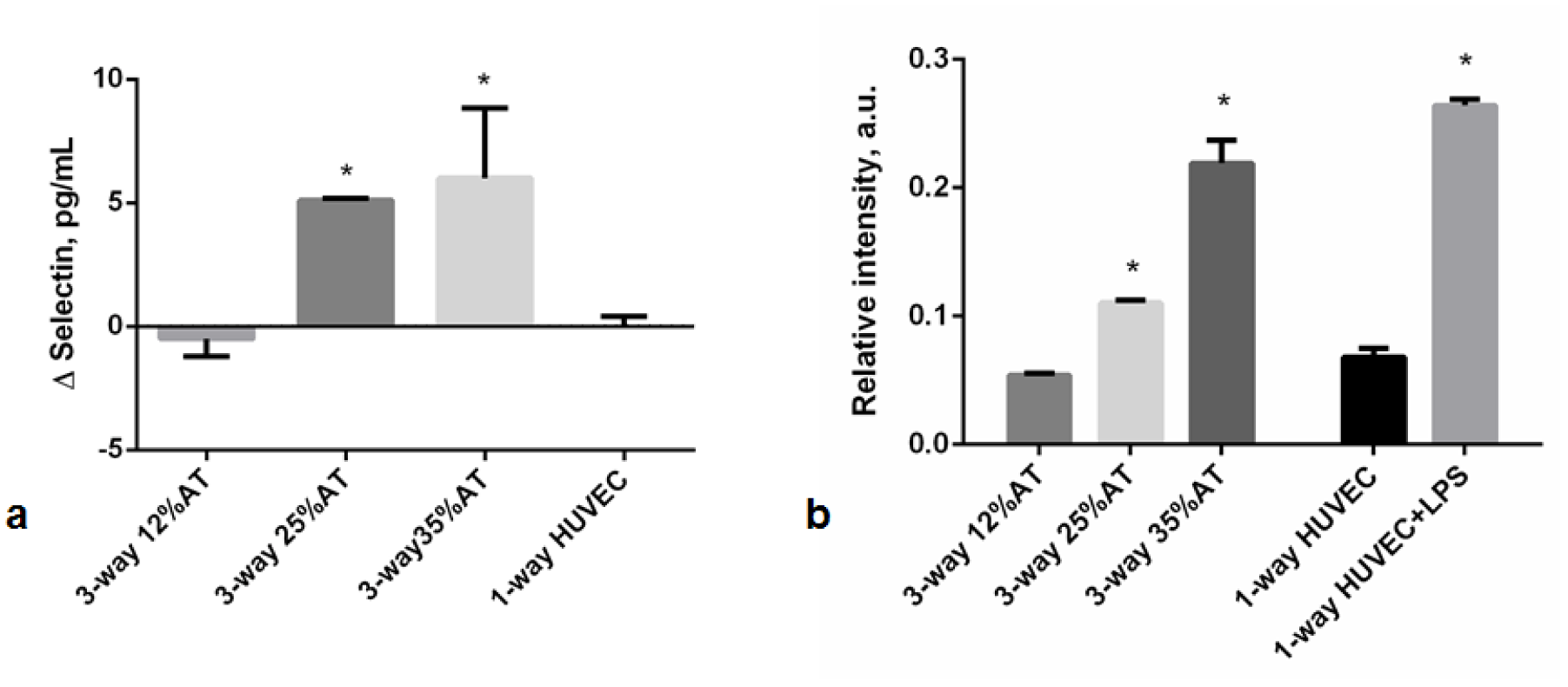
Endothelial-specific markers at 24h. a) Changes in E-selectin concentration in 3-way connected cultures and in HUVEC-1-way control. *= p<0.0001 with respect to control; b) Relative vWF fluorescence intensity in 3-way connected cultures and HUVEC-1-way controls. *=p<0.05 with respect to the corresponding LPS-free 1-way control (n=3 independent experiments for data each point).

## 4. Discussion

Hepatocytes seeded on 3D porous collagen scaffolds, endothelial cells and adipose tissue were cultured together in a fluidic system. Keeping the hepatic and endothelial compartments constant, the quantity of adipose tissue (AT) was varied to represent central or visceral obesity (35%AT), overweight (25%AT) and normo-weight (12%AT) respectively. Our aim was to investigate the interactions among the three tissues as a function of the percentage of adipose tissue as a first step towards the development of a non-animal model for studying the effects of obesity in humans. Given the association between adiposity and hyperlipidemia as well as the causal link between obesity and systemic inflammation [32], we focused primarily on lipid-related molecules and pro-inflammatory markers, examining the similarities and differences between the 3-way model and evidence from human studies in the literature.

All our experiments were conducted with normal glucose medium concentration (5.5 mM), corresponding to a fasting state. Under the fasting state, adipose tissue provides free fatty acid (FFA) and glycerol to the energy expenditure organs by hydrolysing triglycerides. FFAs are also released when adipose tissue is in excess and transported through the vascular system by albumin [33]. They are considered an important link between obesity, insulin resistance, inflammation, development of Type 2 diabetes and associated cardiovascular disorders [34]. In vivo, the elevated plasma FFA levels impair endothelial function [35] and are also associated with an increase in vascular stress markers such as E-selectin [36]. In both the 1-, 2- and 3-way connections, triglycerides were released into the medium, but their levels were significantly higher in the presence of increased adiposity (35%AT and 25%AT) in the 3-way circuit, suggesting that the presence of other tissues stimulated their production in the adipose tissue (Fig 3a). However, in our experiments, FFA levels did not correlate with adiposity in the 3-way system, but were identical for 12%AT, 25%AT and 35%AT. The fairly stable (compared with 1- and 2-way circuits) FFA levels were associated with a significant increase in human albumin concentration – this may be due to the fact that any excess FFA in the 3-way 35%AT and 25%AT conditions was bound to albumin, and hence not detectable in the media. These results are coherent with in-vivo findings, as several studies on humans report a positive correlation between elevated serum albumin levels, nutritional status and body mass index (BMI), likely associated with albumin’s role as a carrier of FFAs [37,38].

Adipose tissue is an important source of inflammatory cytokine production and obesity is regarded as a state of chronic, low-grade inflammation [3]. One of the principal cytokines, interleukin-6 (IL-6), is involved in inflammation and infection responses as well as in the regulation of metabolic, regenerative, and neural processes. It has been observed that adipocytes from obese individuals secrete IL-6, an activity which is correlated with adipocyte volume [39]. MCP-1 is a small inducible cytokine that belongs to the CC chemokine family which recruits monocytes, memory T cells, and dendritic cells to the sites of inflammation produced by either tissue injury or infection [40,41]. Shindu et al. showed that human obesity is associated with an elevated IL-6 expression in the adipose tissue, with increased tissue expression of TNF-α, MCP-1, IP-10 and infiltration by macrophages as the underlying features of a chronic low-grade inflammation [42]. They also found that the MCP-1 upregulation in obese adipose tissue samples correlated with increased IL-6 and suggested that the latter induces subsequent metabolic inflammation.

IL-6 concentrations increased significantly when HUVECs and hepatocytes were connected to 35%AT or 25%AT, compared with the 1-way and 2-way controls and 3-way 12%AT. The increased IL-6 levels were not due simply to the increased proportion of AT, but also a consequence of the cross-talk between the 3 tissue types as verified by comparing the cytokine concentrations in the 3-way connections with those in the presence of 1- or 2-way 12%AT, 25%AT and 35%AT (Fig. 4a). In fact, the interaction analysis shows that the 3-way cross-talk has a substantial upregulating effect (p<0.0001) on the expression of IL-6 in relation to adiposity. In the case of MCP-1 expression, the interaction effect was not significant as we observed an increase in MCP-1 with adiposity in both 1-, 2- and 3-way groups (Fig 4b). However, the increase in the obese (35%AT) 3-way group was significant compared with all other 3-way conditions (p<0.0183, 2-way ANOVA). Similarly, the MCP-1 levels in the 3-way 35%AT were significantly higher than all other obese conditions (ie 1-way 35%AT and 35%At 2-way, see SI). These data suggest that, in our context, MCP-1 induction is closely linked to the amount of adipose tissue and the degree of induction may be stimulated by IL-6, similar to Shindu et al.’s findings in humans [42].

A marked dependency on adiposity was also observed for the endothelial specific inflammatory markers. In the 12%AT 3-way connection, medium E-selectin levels were very low and remained stable over time, while high levels were released into the medium in 35%AT and 25%AT 3-way systems. Similar results were observed for vWF expression, the only difference being the much higher levels of vWF fluorescence in the 3-way 25%AT with respect to 35%AT, 12%AT and 1-way HUVEC controls. Together, these observations indicate that an increasing amount of adipose tissue may cause early endothelial damage and vascular stress, as already demonstrated in-vivo, in humans [43,44].

Obesity is a complex condition which is characterised by an increase in the number of adipocytes as well as the amount of fat stored in the cells [45]. Moreover, obesity is also related to infiltration of adipose tissue by macrophages [46]. Indeed, adipose tissue consists of a variety of different cell types: adipocytes, preadipocytes, stromal/vascular cells and macrophages [47]. We used adipose tissue from donors who did not suffer from excess weight or metabolic disorders, in order to minimise variability due to differences in cell volumes or increased endogenous inflammation which may arise in obese, overweight or diabetic samples. Therefore, in our models, the normal, overweight, and obese state in both 1 and 3-way connections were simulated using increased amounts of adipose tissue and the adipose cell size remained unchanged before and after the experiments over 24 hours.

Although the capacity of HepG2 cells to metabolize xenobiotcs is limited, this immortalized human hepatocyte cell line is generally considered a suitable model of endogenous metabolism [48]. For this reason, although freshly isolated hepatocytes would be more physiological and likely more predictive, we preferred to use the stable HepG2 cell line rather than add a new donor-dependent variable to the system. However, the use of human hepatocytes and long term, chronic experiments which also consider changes in adipose cell volume over time are certainly amongst the next steps to consider for improving the system to make it more physiologically relevant. Additionally, more extensive metabolic and inflammatory profiling of the models would help contribute to a better understanding of the cross-talk between the tissues and strengthen their predictive potential.

In conclusion, the data show that an increase of adiposity in-vitro determines a pro-inflammatory state and results in endothelial stress as observed in vivo. The presence of the 3 tissues in the system is crucial to obtain the integrated, synergic inflammatory responses observed. This first example of an in-vitro model of central obesity using interconnected tissue models clearly demonstrates that it is possible to recapitulate the main features of adiposity related systemic and cardiovascular stress typically observed in humans without resorting to animal models. Engineered non-animal models, in which multiple tissue interactions can be studied piecewise, may help unravel some of the mechanisms of obesity induced metabolic disorders and could, in the future, aid in the design of novel therapeutic interventions. Our study demonstrates the feasibility of this approach and paves the way for further investigations focusing on chronic obesity, lipo-rich diets, hyperglycaemia, and patient-specific responses.

## Acknowledgements

The work leading to these results has received funding from the European Union Seventh Framework Programme (FP7/2007-2013) under grant agreement 304961 (ReLiver). The funders had no role in study design, data collection and analysis, decision to publish, or preparation of the manuscript.

## SUPPLEMENTARY INFORMATION

Figures S1, S2 and S3 show the data for all metabolites and markers at 4 and 24 hours. In the main text we focused on the 24 hour data in order to highlight changes as well as for the sake of comparison with our earlier study. In general, the metabolites were more stable than the pro-inflammatory markers, which tended to increase from 4 to 24 hours. Figure S4 reports the whole dataset showing changes in IL-6 and MCP-1 levels at 24 h for all the 11 conditions (1-way HUVEC, 1-way Hepatocytes and the 9 AT dependent 1-, 2- and 3-way conditions. The Figure also shows the 2 way ANOVA analysis for MCP-1 and IL-6 focusing on the way effect (the AT affect is reported in the main text).

**Figure S1:**
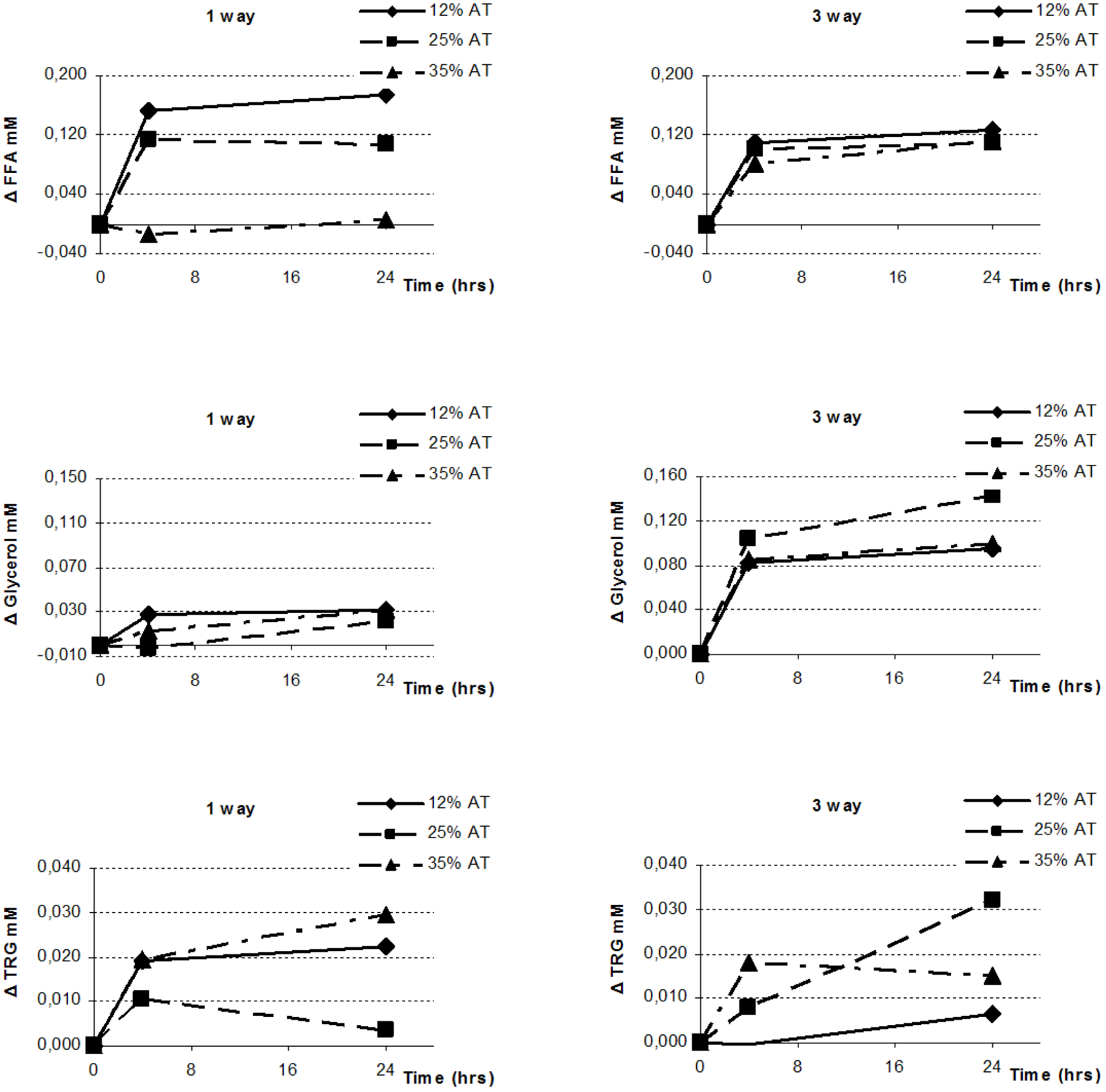
Changes in FFA, glycerol, and triglyceride concentrations in the 1 and 3-way cultures with different amounts of adipose tissue and at different time points (4 and 24h). Metabolite medium concentrations at time 0 were subtracted from to the medium concentrations at various time points.

**Figure S2:**
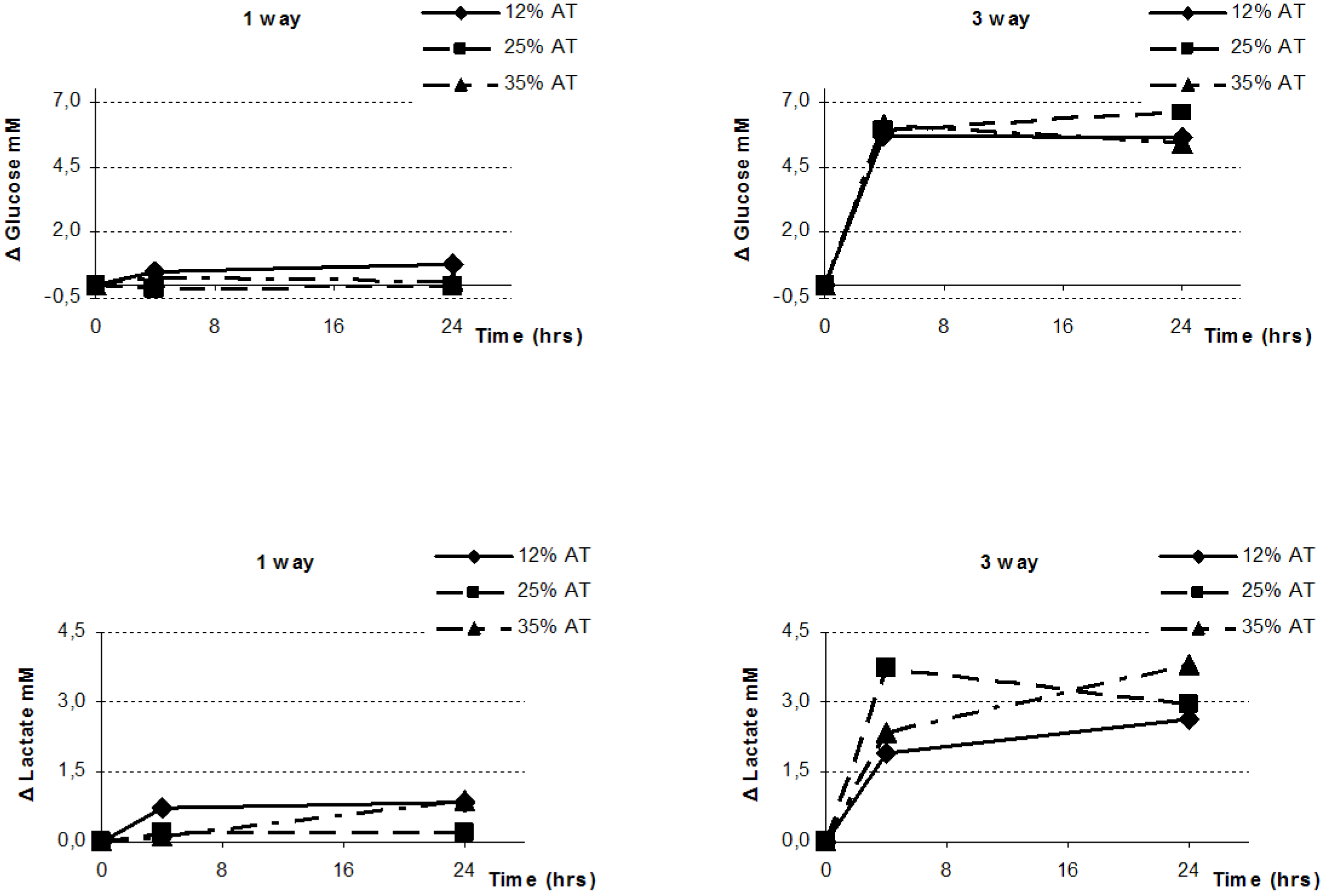
Changes in glucose and lactate concentrations in the 1 and 3-way cultures with different amounts of adipose tissue and at different time points (4 and 24h). Metabolite medium concentrations at time 0 were subtracted from to the medium concentrations at various time points.

**Figure S3:**
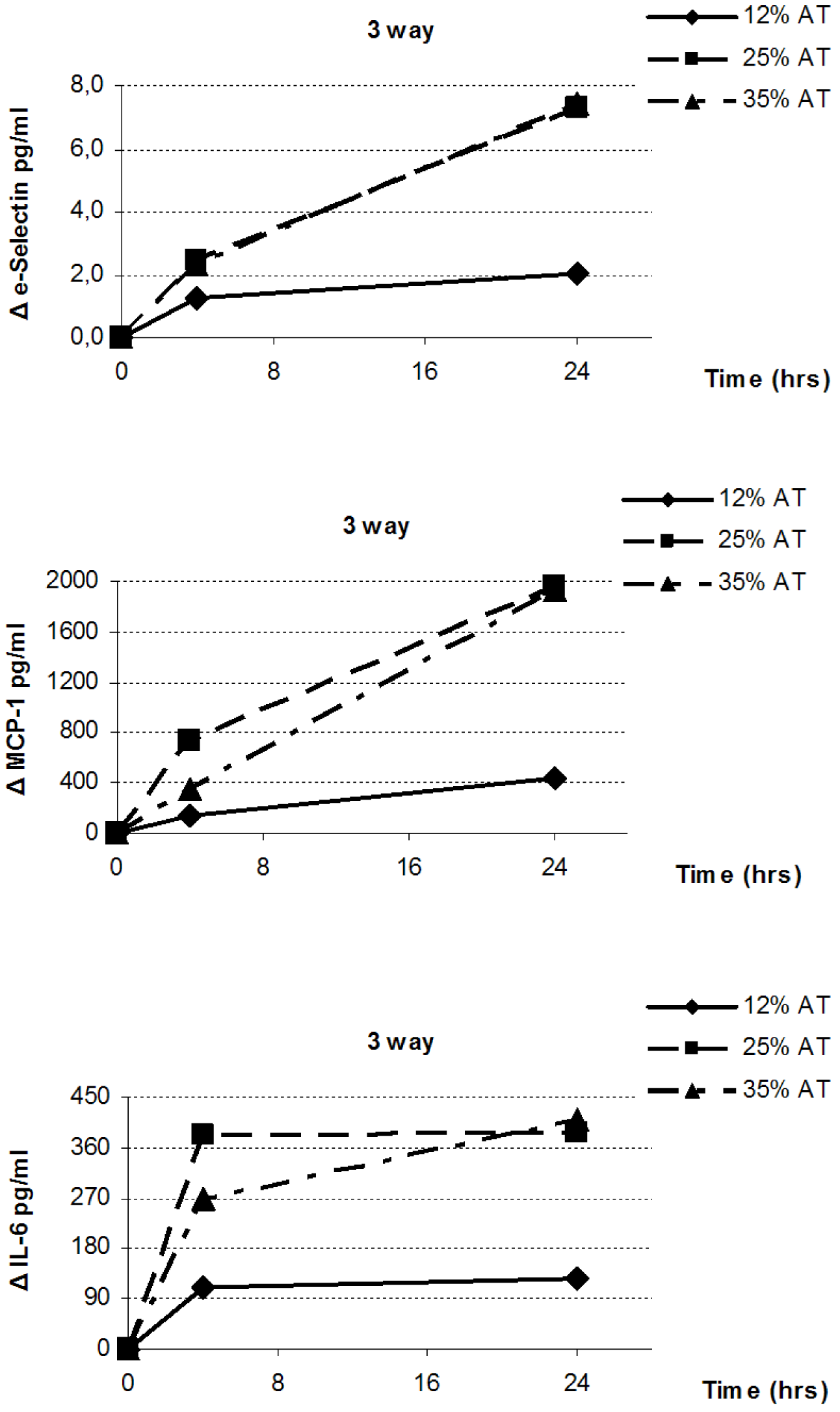
Changes in IL-6, MCP-1 and e-selection concentrations in the 3-way cultures with different amounts of adipose tissue and at different time points (4 and 24h). The concentrations at time 0 were subtracted from the medium concentrations at various time points.

**Figure S4:**
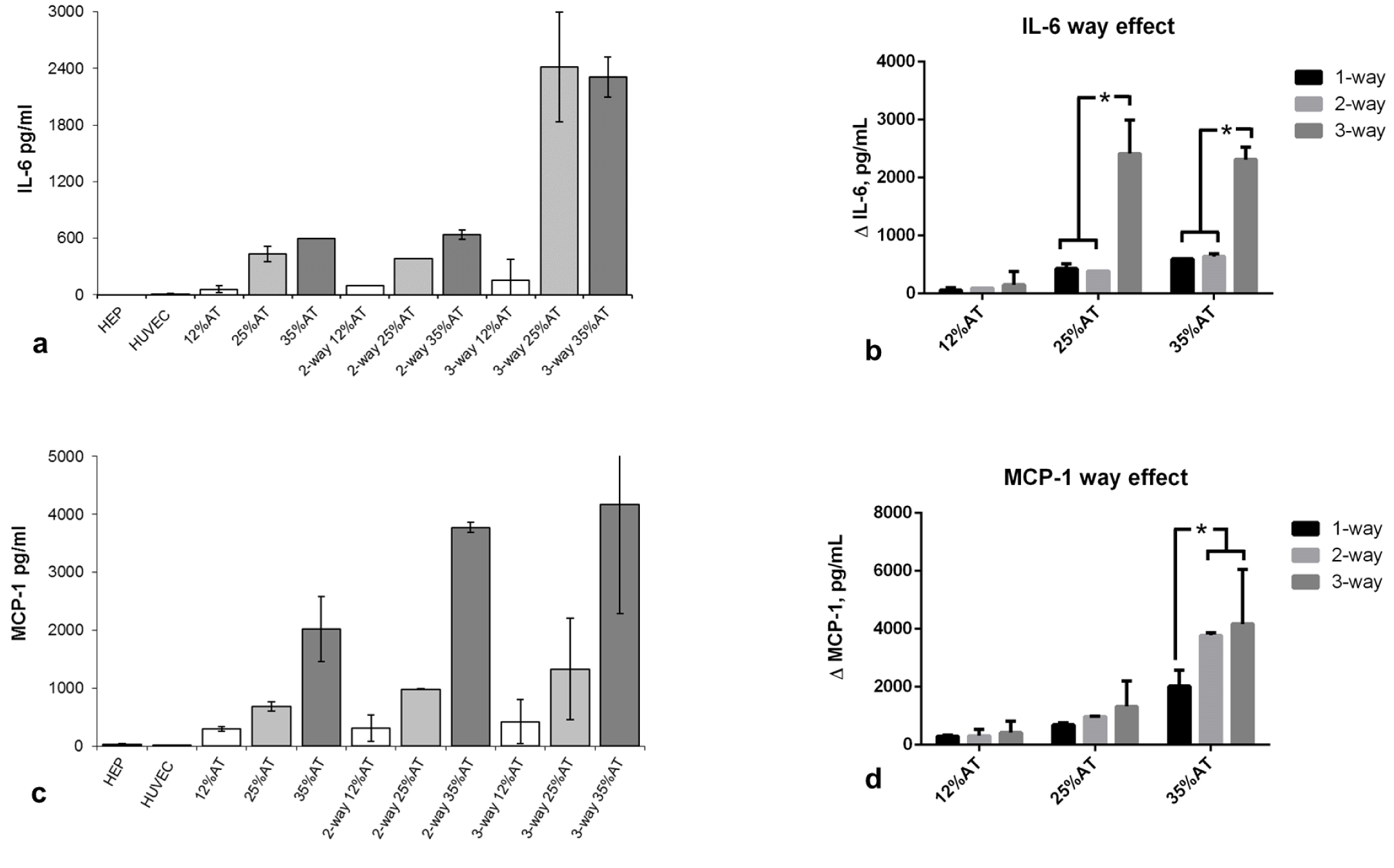
IL-6 and MCP-1 a) IL-6 concentrations at 24h in all conditions; b) 2-way ANOVA analysis for IL-6 showing the way-effect; c) MCP-1 concentrations at 24h in all conditions; c) 2-way ANOVA for MCP-1 showing the way effect. *=p<0.05 In b and c, the concentrations at time 0 were subtracted from the medium concentrations at various time points.

In addition to free fatty acids, glycerol, triglycerides and pro-inflammatory markers, we also measured glucose and lactate. Both metabolites were detected with the Yellow Springs Glucose 2300 STAT (Analytical Service srl, Cassina de Pecchi-MI,Italy).

### Glucose

Although the experiments were conducted using media with 5.5 mM glucose, representing an initial fasting state, in the 1-way adipose controls glucose levels remained fairly stable over 24 hours, suggesting that a different font of energy was used by the these cells. In the 3-way connection, where the total number of cells was higher, a significant increase of glucose consumption was observed, particularly in the presence of 12%AT and 35%AT. In the 25%AT 3-way configuration glucose uptake was reduced with respect to the other 3-way conditions and medium levels remained stable over 24 hours. The 2-way ANOVA analysis shows significant interaction between the tissue cross-talk and adiposity (p=0.004). Glucose uptake in the 1-way connection was almost negligible for all levels of adiposity. As shown in Figure S5, glucose uptake in the 3-way system is not correlated with the amount of adipose tissue or the total number of cells but increases significantly in the presence of 12%AT and 35%AT with respect to the 1-way controls (p<0.005).

**Figure S5:**
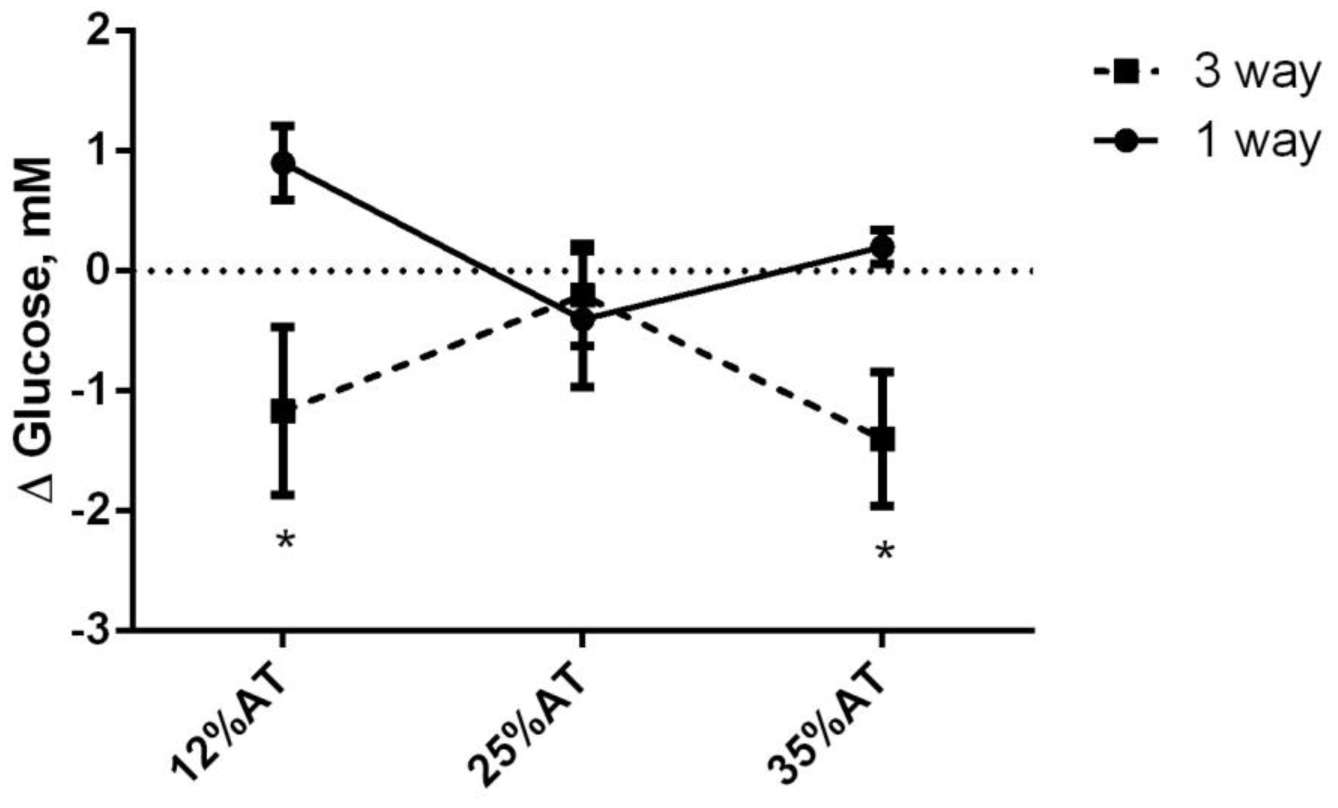
Changes in glucose concentration in control 1-way and 3-way connected cultures as a function of adiposity. * p<0.05 with respect to the corresponding 1-way condition. Lactate

Lactate is considered one of the main glucogenic precursors in the fasting state (*1*) and adipose tissue is an active producer of this metabolite. In-vivo, lactate production is regulated by the nutritional state and the degree of obesity (*2*). Moreover, increased lactate production is observed in adipocytes derived from obese adipose tissue. A net production of lactate was observed in the 3-way connections (Fig. S6, p<0.0001 with respect to the 1-way group, ANOVA2), with the interaction plot indicating the synergic effect of inter-tissue cross-talk on lactate production in the 3-way connection compared with the 1-way adiposity controls. In the 1-way control system medium lactate levels were not correlated with the adipose tissue amounts. However, the presence of endothelial cells and hepatocytes in the 3-way circuit determined a significant increase in lactate levels, mirroring the trend observed in-vivo.

**Figure S6:**
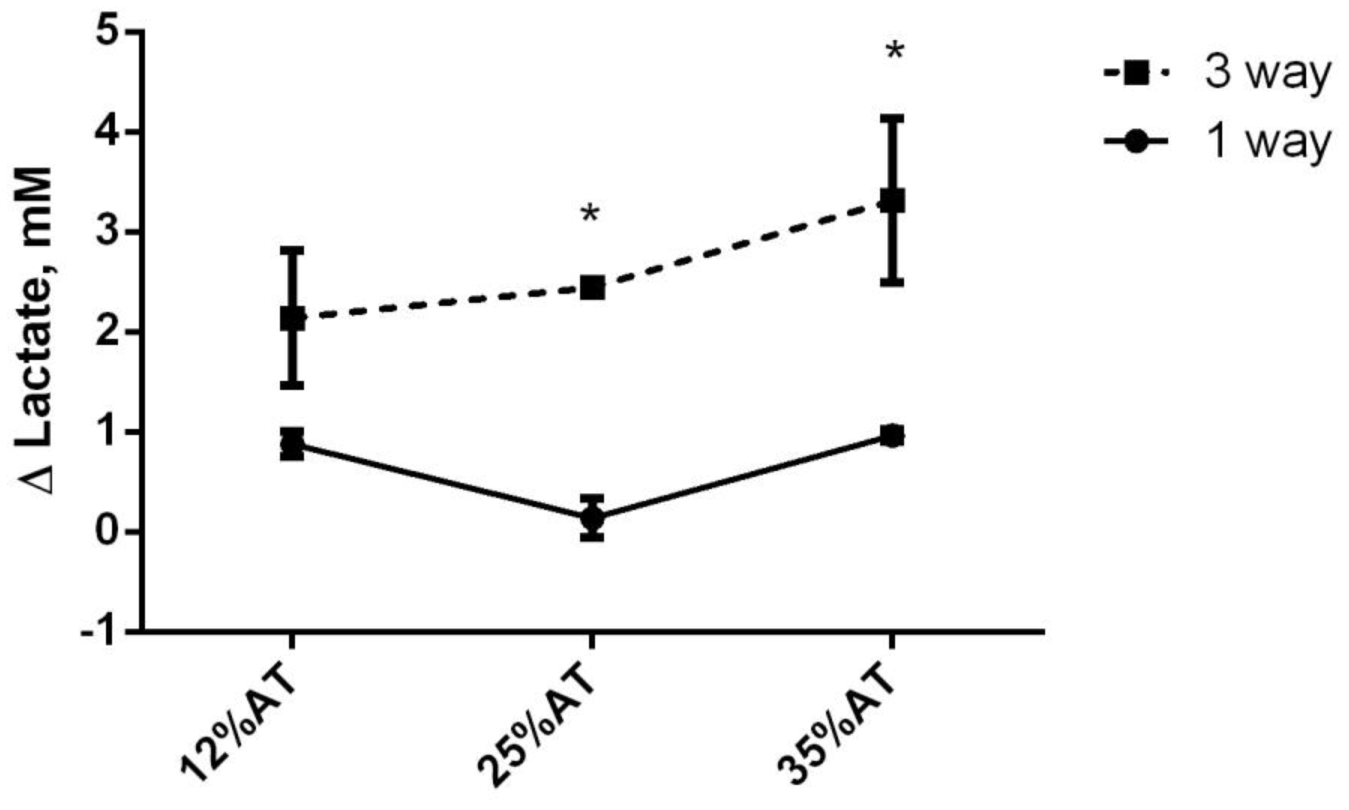
Changes in lactate concentration in control 1-way and 3-way connected cultures as a function of adiposity. Different letters indicate significant differences.

### Estimation of Von Willebrand factor (vWF) expression through image processing

The amount of vWF expressed per endothelial cell in different conditions was measured using immunohistochemistry. At the end of the 24 h experiments, the bioreactor circuit was dissembled by first draining the media reservoir and separating the 3 modules. The laminar flow Ibidi slide with HUVEC was treated with 4% paraformaldehyde for 20 mins as per manufacturer’s instructions and cells were stained for vWF with Monoclonal Mouse Anti-hvWF clone F8/86 (Dako, Denmark) and anti-mouse FITC (Invitrogen, Paisley, UK) as a secondary antibody; DAPI (4’,6-diamidino-2-phenylindole) staining was also performed for nuclear labeling. Fluorescence was visualized using a confocal microscope (Nikon A1). In particular, images were taken using a 10X objective with a pixel-to-micron ratio of 1.23 μm/pixel on a 1024x1024 matrix. For the green channel, the same confocal settings were used for all scans (i.e. 25 W laser power). To quantify vWF expression, firstly, the number of nuclei (#nuclei) was calculated on the blue channel of each image using ImageJ. As regards the green channel, a global threshold using the Otsu method *(3)* was performed, then the Mean Pixel Intensity (MPI) per nucleus (MPI/#nuclei) was evaluated. The MPI was calculated using the method described in Gonzalez et al., 2009. *(4)*.

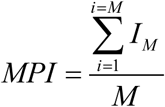

where M is the number of the object pixels (i.e. the cell regions expressing vWF) and Im their pixel intensity.

Data were averaged over 3 samples per condition (1-way HUVEC, 1-way HUVEC+10 ng/mL LPS and 3-way 12%AT, 25%AT and 35%AT) and at least 5 regions of interest per sample. Sample images are reported in Figure S7.

**Figure S7:**
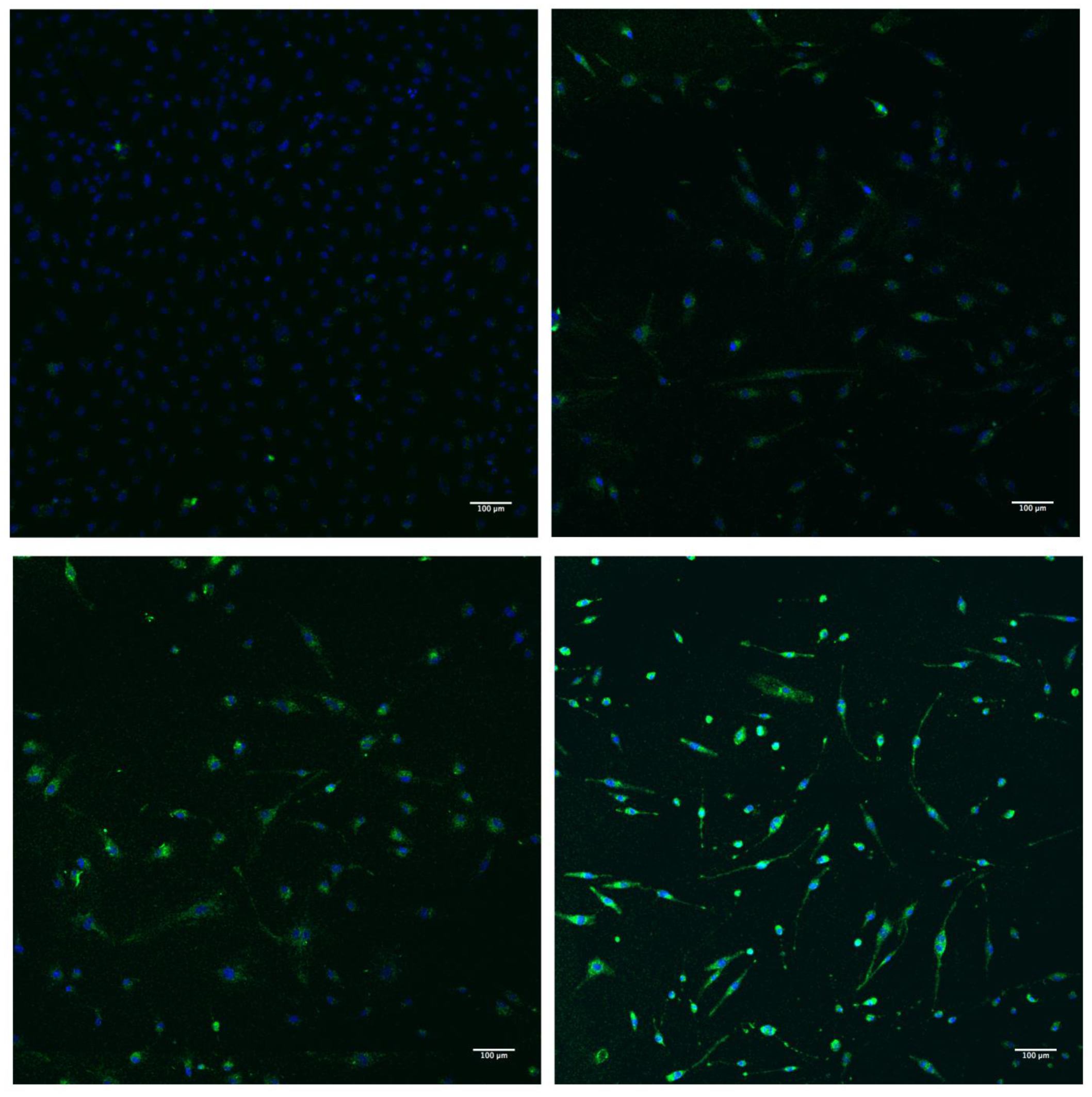
*vwF (green), DAPI (blue) stained HUVEC after 24h. A) HUVEC control, B) 3-way 12%AT, C) 3-way 25%AT, D) 3-way 35%AT.*

